# The chromosome-level genome of the ctenophore *Mnemiopsis leidyi* A. Agassiz, 1865 reveals a unique immune gene repertoire

**DOI:** 10.1101/2024.01.21.574862

**Authors:** Vasiliki Koutsouveli, Montserrat Torres-Oliva, Till Bayer, Janina Fuß, Nora Grossschmidt, Angela M. Marulanda-Gomez, Diana Gill, Ruth A. Schmitz, Lucía Pita, Thorsten B. H. Reusch

**Author notes:** Equal contribution to the manuscript (Vasiliki Koutsouveli, Montserrat Torres-Oliva). **Authors for Correspondence:** Lucía Pita, Institut de Ciències del Mar–Spanish National Research Council (CSIC), Marine Biology and Oceanography, Marine Biogeochemistry, Atmosphere and Climate, Barcelona, Spain, Thorsten B. H. Reusch, Division of Marine Ecology, Marine Evolutionary Ecology, GEOMAR Helmholtz Centre for Ocean Research Kiel, Kiel, Germany, TBHR.

## Abstract

Ctenophora are basal marine metazoans, the sister group of all other animals. *Mnemiopsis leidyi* is one of the most successful invasive species worldwide with intense ecological and evolutionary research interest. Here, we generated a chromosome-level genome assembly of *M. leidyi* with a focus on its immune gene repertoire. The genome was 247.97 Mb, with N50 16.84 Mb, and 84.7% completeness. Its karyotype was 13 chromosomes. In this genome and that of two other ctenophores, *Bolinopsis microptera* and *Hormiphora californensis*, we detected a high number of protein domains related to potential immune receptors. Among those, proteins containing Toll/interleukin-1(TIR2) domain, NACHT domain, Scavenger Receptor Cystein-Rich (SRCR) domain, or C-type Lectin domain (CTLD) were abundant and presented unique domain architectures in *M. leidyi. M. leidyi* seems to lack *bona fide* Toll like Receptors, but it does possess a repertoire of 15 TIR2-domain containing genes. Besides, we detected a *bona fide* NOD-like receptor and 38 NACHT-domain containing genes. In order to verify the function of those domain containing genes, we exposed *M. leidyi* to the pathogen *Vibrio coralliilyticus*. Among the differentially expressed genes, we identified potential immune receptors, including four TIR2-domain containing genes, all of which were upregulated in response to pathogen exposure. To conclude, many common immune receptor domains, highly conserved across metazoans, are already present in Ctenophora. These domains have large expansions and unique architectures in *M. leidyi*, findings consistent with the basal evolutionary position of this group, but still might have conserved functions in immunity and host-microbe interaction.

## Introduction

Ctenophora, or comb jellies, are a unique phylum of exclusively marine invertebrates. They feature a mostly pelagic life-style as zooplankton feeders (only very few benthic species exist), and are distributed across all oceans, temperature gradients and depths (Tamm 2014; Dunn et al. 2015). Ctenophores are the earliest diverging phyla in the evolution of metazoans, containing around 200 described species and likely many more undescribed to date (Tamm 2014; Dunn et al. 2015). The lively debate on whether or not sponges or comb jellies appeared first in the tree of life (Ryan et al. 2013; Moroz et al. 2014; Halanych 2015; Whelan et al. 2015; Pisani et al. 2015; Feuda et al. 2017; Kapli & Telford 2020) was recently settled. Schultz et al. (2023) demonstrated that ctenophores are the sister group to all other metazoan phyla based on chromosomal synteny. This is in line with many other comparative studies of the last decade that revealed that this phylum is unique in terms of many traits such as neural system and locomotion as well as cell type organisation (Martindale et al. 2002; Jager et al. 2010; Moroz et al. 2014). Up to date, studies in ctenophores have shed light on the evolutionary origins of bioluminescence, the evolution of the neural system and muscular organization, the Wnt signalling, axis patterning and locomotion, as also wound healing and adult regeneration (Martindale et al. 2002; Schnitzler et al. 2012; Jager et al. 2013; Moroz 2015; Jager & Manuel 2016; Babonis et al. 2018; Ramon-Mateu et al. 2019; Burkhardt 2022). In contrast, only few studies have focused on the immunological repertoire of ctenophores (Bolte et al. 2013; Moroz et al. 2014; Traylor-Knowles et al. 2019), while, the innate immune system in other non-bilaterian phyla, such as sponges and cnidaria, showed an impressively high molecular complexity(Wiens et al. 2007; Augustin & Bosch 2010; Srivastava et al. 2010; Detournay et al. 2012; Bosch 2013, 2014; Hentschel et al. 2012; Krishnan et al. 2014; Riesgo et al. 2014; Degnan 2015; Pita et al. 2018; Parisi et al. 2020; Emery et al. 2021; Schmittmann et al. 2021). This fact leads to the necessity to study further the immune repertoire in ctenophores, expecting to discover similar complexity and unique features in this group.

In the animal world, hosts live in an environment full of microbes in which they have evolved in symbioses ranging from reciprocal benefits to antagonistic relationships. Under this scenario, the animal innate immune system acts not only as defence mechanism against microbial infections but it is also key for maintaining and specifying interkingdom communication between host and their microbiota (Koropatnick et al. 2004; Nyholm & Graf 2012; Bosch 2013; Chu & Mazmanian 2013; Gerardo et al. 2020). Like other aquatic organisms, ctenophores have established mutualistic relationships with microbial communities (Daniels & Breitbart 2012; Breitbart et al. 2015; Jaspers et al. 2019), evolving together as holobionts (Bosch & McFall-Ngai 2011). As such, it is expected that ctenophores already possess developed immune mechanisms to control their microbiota and distinguish between mutualists and pathogens. A crucial first step in animal-microbe interaction is the recognition of Microbial-Associated Molecular Patterns (MAMPs), such as lipopolysaccharides (LPSs), peptidoglycan (PGN), and flagellin, by host Pattern-Recognition Receptors (PPRs), which further trigger downstream pathways including several transcription factors and modulators (Zheng et al. 2005). There is a large variety of PRRs among animals, of which the most common ones are the Toll-Like Receptors (TLRs), the Nucleotide Oligomerization Domain (NOD)-Like Receptors (NLRs), the C-Type Lectins (CTL) and the scavenger receptors (SRs) (Rosenstiel et al. 2009). Apart from the typical PPRs, other receptors which have been related to microbial recognition are the G Protein-Coupled Receptors (GPCRs), and cytokine receptors (Reboul & Ewbank 2016; Liongue et al. 2016). Most of these receptors are characterized by specific protein domains and motifs (Dierking & Pita 2020). In marine invertebrates, these receptors are highly divergent, and they can be either present fully or with parts of their original architecture, containing some of the homologous domains’ architecture (e.g. Riesgo et al. 2014). Previously, it has been reported that the ctenophore *Pleurobrachia bachei* lacks the common pattern recognition receptors and further mediators of the innate immune system (e.g. TLRs and TIR domain, Myd88, NLRs, RLRs, etc) found in all other bilaterians and sponges (Moroz et al. 2014), whereas the ctenophores *Mnemiopsis leidyi* and *Hormiphora californensis* possess several of these conservative genes or their related domains (Traylor-Knowles et al. 2019). Consequently, it becomes crucial to identify the immune repertoire in ctenophores in order to extend deeper our understanding on their biology as well as the origin and evolution of the innate immune system in metazoans.

Chromosome-level genomes have become indispensable to characterize the gene repertoire of non-model (marine) species (e.g., Kenny et al. 2020; Cazet et al. 2023). So far, the majority of chromosome-scale genome assemblies come from bilaterians, and only a low number of chromosome-level genome assemblies is currently available from nonbilaterian animals, mainly from cnidarians (Zimmermann et al. 2020; Li et al. 2020; Kon-Nanjo et al. 2023; Cazet et al. 2023) and the freshwater sponge *Ephydatia muelleri* (Kenny et al. 2020). Regarding the Ctenophora phylum, two chromosome-level genomes have been recently published from the species *Hormiphora californensis* and *Bolinopsis microptera* (Schultz et al. 2021, 2023).

Here, we generated a chromosome-level assembly of the widespread ctenophore species *M. leidyi* Agassiz, 1865 with Pacbio and Hi-C technologies, in order to further explore the immune repertoire in this animal. *M. leidyi* is native to the western Atlantic Ocean, but in the last decades it spread as one of the most invasive species to many European waters and to the western Asian region (Reusch et al. 2010). Its notorious expansion in the Black Sea, the Caspian Sea and North and Baltic Seas was associated with a decrease in the fish population due to fish larvae predation, and caused huge ecological and economic impact. Moreover, this species also became an important model species for basic biological research, in particular in the fields of bioluminescence (Schnitzler et al. 2012), body regeneration (Henry & Martindale 2000; Ramon-Mateu et al. 2019), axis patterning (Martindale et al. 2002), and population genetics (Ghabooli et al. 2011; Bolte et al. 2013; Verwimp et al. 2020; Jaspers et al. 2021).

Furthermore, its easy collection in the field, the establishment and maintenance of cell cultures, and its successful reproduction *in vitro* have turned *M. leidyi* into a model species for evolutionary and developmental studies (Martindale 2022). Hence, obtaining a chromosome-scale genome will also provide crucial genomic resources for many other studies, along with our analysis on the immune gene repertoire.

## Results

### *M. leidyi* chromosome-level genome assembly

High molecular weight DNA (total amount of 726.66 ng/μl; nanodrop A260/280:1.87; A230/260: 2.49) was extracted from one individual of *M. leidyi* collected in the Kiel Fjord (Figure 1A). A draft assembly was initially generated using a PacBio Sequel II (PacBio, Menlo Park, CA). Data from two SMRT cells were collected, generating 728,870 CCS/HiFi reads and a total of 9.2 Gb (∼ 37x coverage based on the final assembly size). Final assembly size of this draft assembly was 247.94 Mb, with N50 1.01 Mb. After mapping the Omni-C libraries to the draft genome assembly, a high-quality chromosome-level genome assembly was generated. This assembly contained 142 sequences and, based on length distribution and Hi-C contact maps, 13 chromosomes were identified (Figure 1B, Supplementary Figure 1). Due to the chromosome-level conservation between ctenophores, we have labelled the 13 assembled chromosomes of *M. leidyi* according to the nomenclature introduced for other two ctenophore species (*B. microptera* and *H. californiensis*, see Supplementary Figure 2 for details). From the 129 scaffold sequences that were not assigned to a chromosome, 127 had a high similarity blast hit (E< 1e-30) to one of the assembled chromosomes, indicating they are alternative haplotype regions. The remaining 2 sequences were mapped against NCBI nr database and shown to be a *Marinomonas* chromosome (CP025532.1 *Marinomonas* sp. A3A chromosome) and a tandem repeat (TACATGGAGTTACAATGT). Therefore, all 129 unassigned scaffolds were discarded from the chromosome-level assembly, and all the following analyses were performed on this clean assembly. This final assembly had a total size of 247.97 Mb, with a chromosome N50 of 16.84 Mb and 39.1% GC content (Table 1). BUSCO scores indicated that 84.7% of single-copy metazoan orthologs were complete in the *M. leidyi* chromosome-level assembly (Figure 1C). After automatic annotation using the *genomeannotator* pipeline (https://github.com/marchoeppner/genomeannotator), a total of 18,972 protein coding genes were identified in the *M. leidyi* assembly (Supplementary Table 1). After functional annotations of this gene set, KEGG pathways were assigned to 5,602 genes, GO terms to 7,404 genes and PFAM domains to 13,282 genes (Supplementary Table 1). BUSCO scores showed that this annotation contains 76.2% complete single-copy metazoan orthologs (Figure 1C).

**Figure 1.**
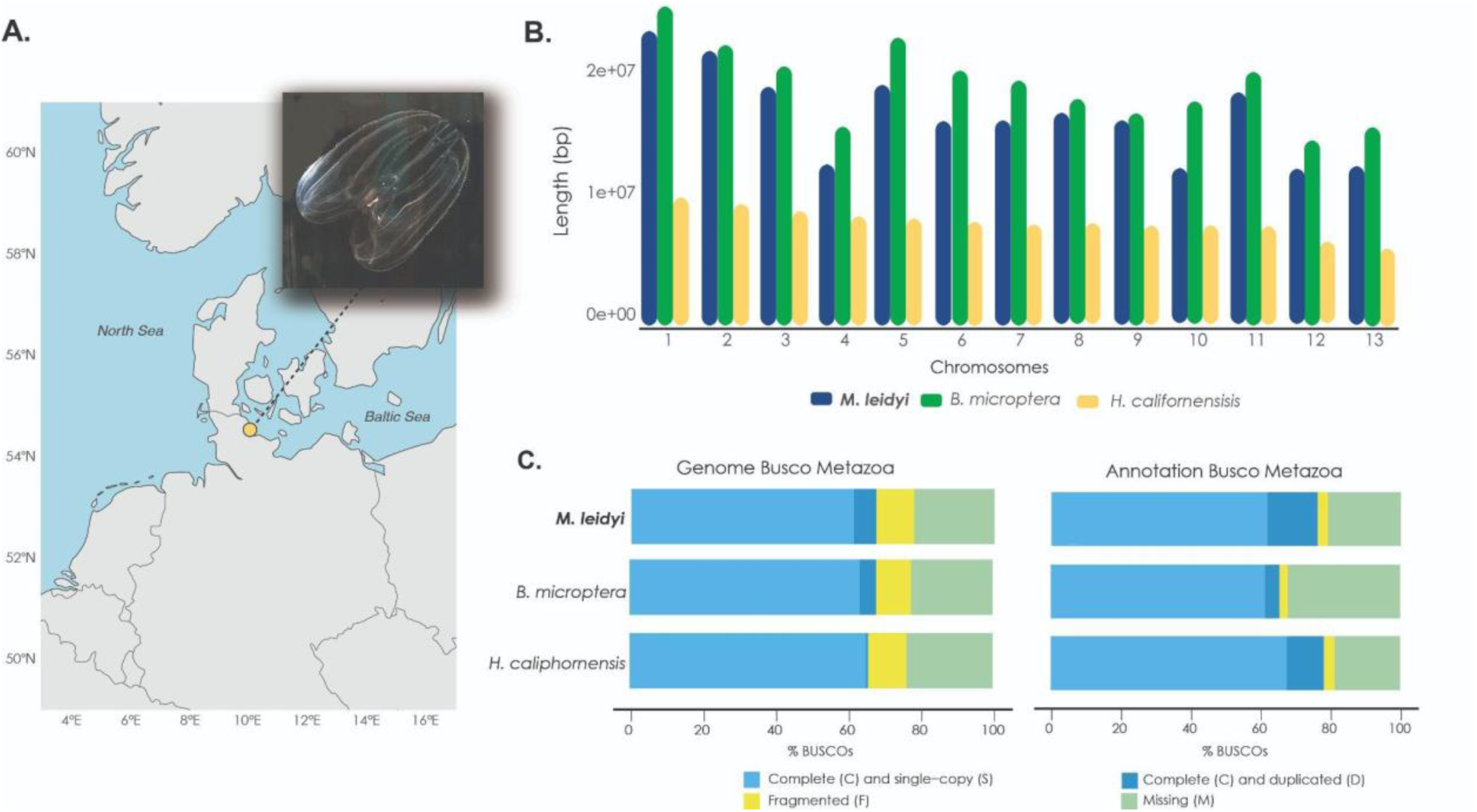
Comparative genome statistics. **A.** Sampling location of the individual *Mnemiopsis leidyi* used for genome sequencing and assembly (depicted in the image)**. B.** Length of the 13 chromosomes found in the genome of *M. leidyi* compared to the analogous 13 chromosomes found in the published genomes of genome of *Bolinopsis microptera* and *Hormiphora californensis*. **C.** Comparison of Busco scores indicating the quality of the three genomes and their annotation. Busco scores were run for each genome and their annotation. The Busco analysis was run with Busco v.5 version with metazoa_odb10 database.

**Table 1.** Comparison of statistics between the chromosome-level genome assemblies of the ctenophore species: *Mnemiopsis leidyi*, *Hormiphora californensis* and *Bolinopsis microptera*.

|  | *Mnemiopsis*<br>*leidyi* | *Bolinopsis*<br>*microptera* | *Hormiphora*<br>*californensis* | *Mnemiopsis*<br>*leidyi* |
| --- | --- | --- | --- | --- |
| **reference** | This study | Schultz et al., 2023 | Schultz et al., 2021 | Ryan et al., 2013 |
| **genome size** | 248 Mb | 265 Mb | 111 Mb | 156 Mb |
| **Number of scaffolds** | 142 | 346 | 44 | 5100 |
| **Number of chromosomes** | 13 | 13 | 13 | Not detectable |
| **Largest chromosome** | 20.0 Mb | 26.2 Mb | 10.5 Mb | na |
| **Number of Ns** | 33.3 kb | 21.1 kb | 30.6 kb | 5.5 Mb |
| **GC content** | 39.1 % | 38.5% | 43.1 % | 37.5% |
| **N50** | 16.8 Mb | 20.8 Mb | 8.5 Mb | 187 kb |

### Search of protein domains common in animal immune receptors

Upon genome annotation, we focused our search on 23 Pfam domains that are known to be part of conserved animal immune receptors (Buckley & Rast 2011). First, we looked for the presence of these domains in the genome of *M. leidyi* and two other ctenophore species, *B. microptera* and *H. californensis* with available chromosome-level genomes (Figure 2A). Out of 23 Pfam domains, 20 domains were detected in the genome of *M. leidyi*, based on the genome annotation and independent hmmersearch (Figure 2A). Most of them were also detected in the other two species, with exception the SEF/IL-17R (SEFIR) domain that was absent in *H. californensis* but present in *M. leidyi* and *B. microptera,* and the Baculovirus IAP Repeat domain (BIR) domain that was absent in *B. microptera* but detected in *M. leidyi* and *H. californensis* (Figure 2A). Interestingly, we did not detect the Toll/interleukin-1 (TIR) Pfam domain (accession number PF01582) in any of the ctenophore species, while the TIR2 Pfam domain was present (accession number PF13676).

**Figure 2.**
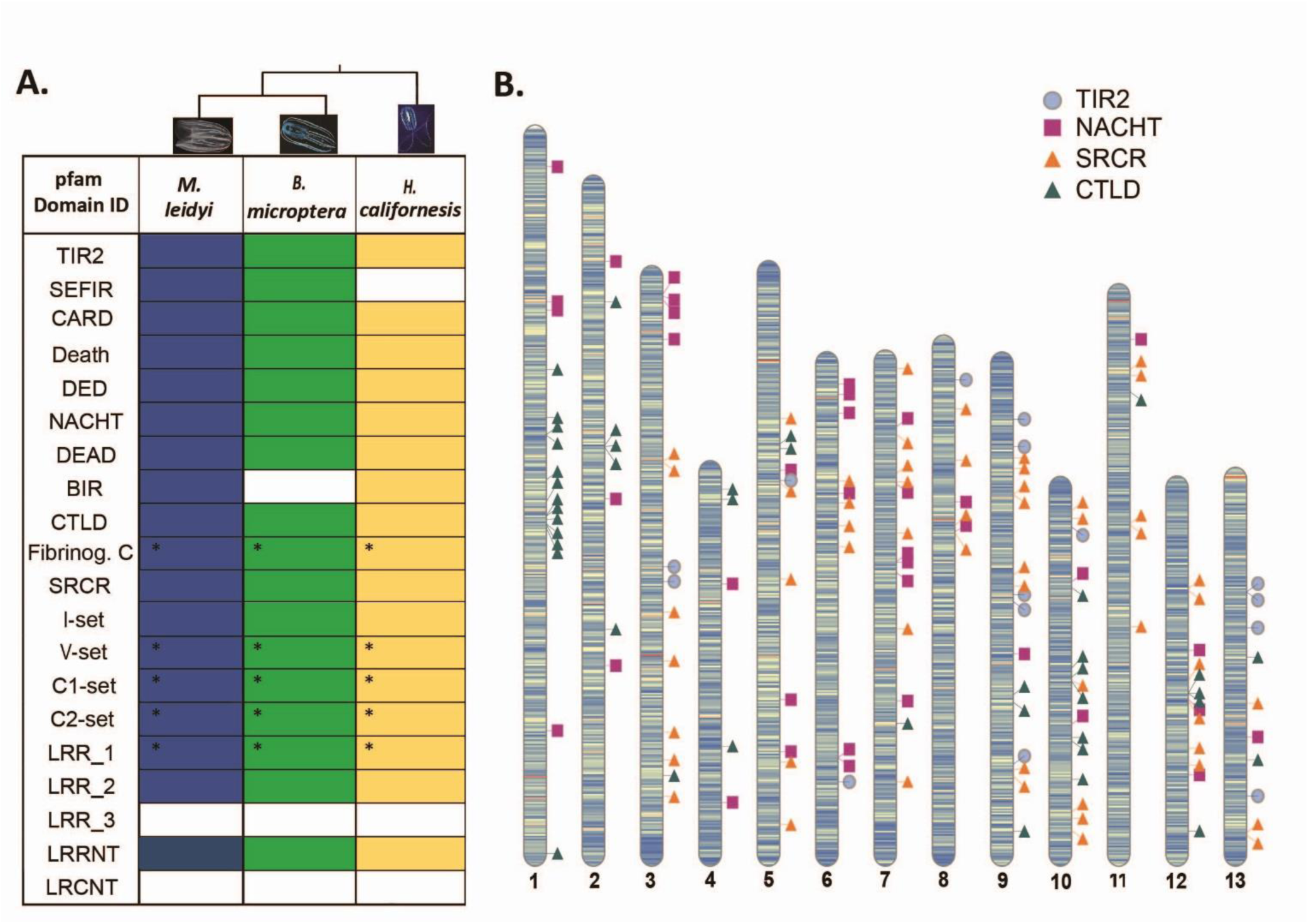
Pfam domains commonly found in immune receptors. **A.** presence/absence of the reported pfam domains in the genome of *Mnemiopsis leidyi*, and the other two ctenophores’ published chromosome genomes, *Bolinopsis microptera* and *Hormiphora caliphornensis*. The evidence for presence or absence of a domain was taken after search in the annotation of the genomes and independent hmmerscan for each domain, based on default parameters. The “*” symbol indicates the pfam domains which were represented by very few sequences with low E-value threshold. Toll/interleukine-2 domain; Immunoglobilin I-set, Immunoglobilin V-set, Immunoglobilin C1-set, Immunoglobilin C2-set, Leucine-rich repeat 1 domain, Leucin-rich repeat 2 domain, Leucin-rich repeat 3 domain, Leucin-rich repeat N-terminal domain, Leucin-rich repeat C-terminal domain, Scavenger Receptor cystein rich domain, Caspase recrutiment domain, DEAD/DEAH box helicase, Death effector domain, Lectin C type domain, Fibrinogen beta and Gamma chains C-terminal globular domain, Inhibitor of Apoptosis domain, 7 transmembrane (TM) helices_1,2,3, **B**. Genes containing the TIR2, NACHT, SRCR, and C-type Lectin pfam domains, are depicted in which chromosome they are located in the genome. These pfam domains were chosen as they are part of the most conserved families of immune receptors in metazoans. The graph was created with RIdeogram.

Out of the above 20 detected domains in the genome of *M. leidyi*, we further focused on four of them because of their importance in the immune repertoire of early metazoans (e.g., Poole and Weis, 2014; Riesgo et al. 2014; Neubauer et al. 2016; Pita et al. 2018; Schmittmann et al. 2021): The TIR2 domain, which is part of the canonical TLRs, the NACHT domain, which is component of the NLRs, the Scavenger Receptor Cystein-Rich domain (SRCR), part of the SRs and the C-type Lectin domain (CTLD) which is part of the CTLs. In total, 153 genes were coding for those domains in the *M. leidyi* genome (Supplementary Table 2). Specifically, 15 genes were coding a TIR2 domain, mainly found in chromosomes 9 and 13 (Figure 2B; Supplementary Table 2), while 114 genes comprised the NACHT domain, spread among different chromosomes with many genes detected in chromosomes 3 and 7 (Figure 2B; Supplementary Table 2). Additionally, 42 genes contained CTLD, 13 of those were in chromosome 1 and 7 genes in chromosome 10. The remaining genes were distributed across all the other chromosomes (Figure 2B; Supplementary Table 2). Finally, 55 genes contained SRCR, most of those detected in chromosomes 7, 9 and 12 (Figure 2B; Supplementary Table 2).

*M. leidyi* proteins containing TIR2, NACHT, SRCR, or CTLD domains revealed a large diversity of architectures (Figure 3). Above domains were combined with a large variety of other domains, motifs and signals, resulting in an overall diverse array of proteins. Among the TIR2 domain containing proteins, canonical TLR based gene models conserved across animals, invertebrates and vertebrates alike, could not be identified. One TIR2 containing protein included the death domain, and might be considered as Myd88-like (Figure 3A). Similarly, among the NACHT containing proteins, a single one contained the LRR domain which putatively could be a *bona fide* NLR, but we were unable to reconstruct the canonical structure of vertebrate NLRs (Figure 3B). Furthermore, we did not find any sequence that corresponds to the NLR receptor in the annotation. The SRCR domain containing proteins had a large expansion in *M. leidyi*. We found proteins with many SRCR repeats and one combined with a CTLD, which, in vertebrates, are common of SR-I and SR-E classes respectively. However, SRCR domains combined with collagen (class SR-A), which is also typical of the vertebrate SRs, were absent. Instead, we found a large number of SRCR-containing sequences with novel and unique domain combinations (Figure 3C). Looking finally at the CTLD containing proteins, they were also highly diverse regarding their overall domain architecture, including combinations with several types of Immunglobulin (IG), Epidermal Growth Factor (EGF), and Sushi domain (CCP) (Figure 3D).

**Figure 3.**
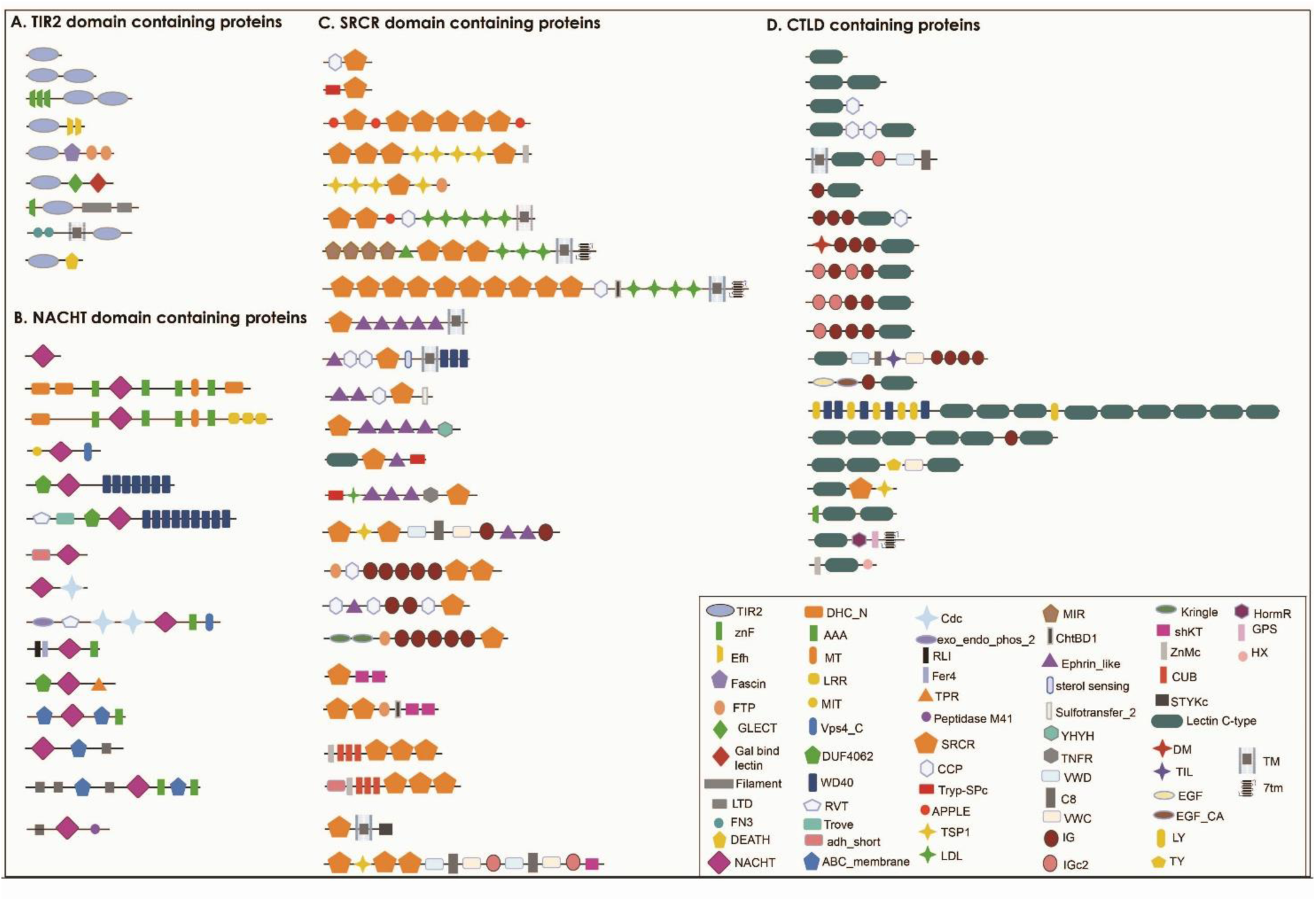
Domain architecture of pfam domain-containing proteins. Domain architecture of proteins potentially comprising the innate immune receptor repertoire of *Mnemiopsis leidyi*, based on the presence of conserved pfam domains **A.** TIR2, Toll/interleukine-1; znF, zink finger domain; Efh, EF hand domain; LTD, C-terminal lamin-tail domain; FN3, fibronectin III. **B.** NACHT, DHC_N, Dynein heavy chain; LRR, leucine-rich repeat; Vps4_c, Vacuolar protein sorting-associated protein 4WD40, repeat of 40 amino acids typically terminating in Trp-Asp; DUF4062, conserved domain of unknown function; RVT, reverse transcriptase; adh_short, short chain dehydrogenase; RLI, metal-binding domain in endoribonuclease RNase L inhibitor; TPR, tetratricopeptide repeat. **C.** SRCR, Scavenger Receptor Cysteine-Rich domain; CCP, complement control protein, known as Sushi; VWD, Von Willebrand factor type D domain, VWC, Von Willebrand factor type C domain, TRyp_SPc, Trypsin-like serine protease; TSP1, Thrombospondin-1; LDL, Low-Density Lipoprotein receptor domain; MIR, named after three of the proteins in which it occurs: protein Mannosyltransferase, Inositol 1,4,5-trisphosphate receptor and Ryanodine receptor; TNFR, tumor necrosis factor receptor; DM, DNA binding domain; IG, Immunoglobulin; IGc2, C2-set immunoglobulin; shKT, a 35-residue peptide toxin from sea anemone Stichodactyla helianthus, are potent inhibitors of K channels; ZnMc, Peptidase metallopeptidase; CUB, complement C1r/C1s, Uegf, Bmp1; STYKc, protein kinase. **D.** CTLD, C-Type Lectin Domain; TIL, trypsin inhibitor-like cysteine rich domain; EGF, epidermal growth factor; EGF_ca, EGF-calcium-binding, epidermal growth factor; GPS, G-protein-coupled receptor proteolytic site; HormR, present in hormone receptors; LY, Ly-6 antigen/uPA receptor-like; TY, Thyroglobulin type I repeats; HX, hemopexin, TM, transmembrane region; 7TM, 7 helix transmembrane domain. The Figure was created with BioRender.com.

The phylogenetic analysis of TIR containing proteins indicated that the TIR2 sequences of ctenophores including *M. leidyi* are closer to TIR sequences of non-metazoans and early diverging metazoans, while they do not possess any TLR-like or Myd88-like proteins (Figure 4). The analysis, including Filasterea, Placozoa, and different phyla of metazoans, divided those proteins into three main groups, one group with only TIR containing proteins, another group with TLR-like proteins and a third group with Myd88-like proteins.

**Figure 4.**
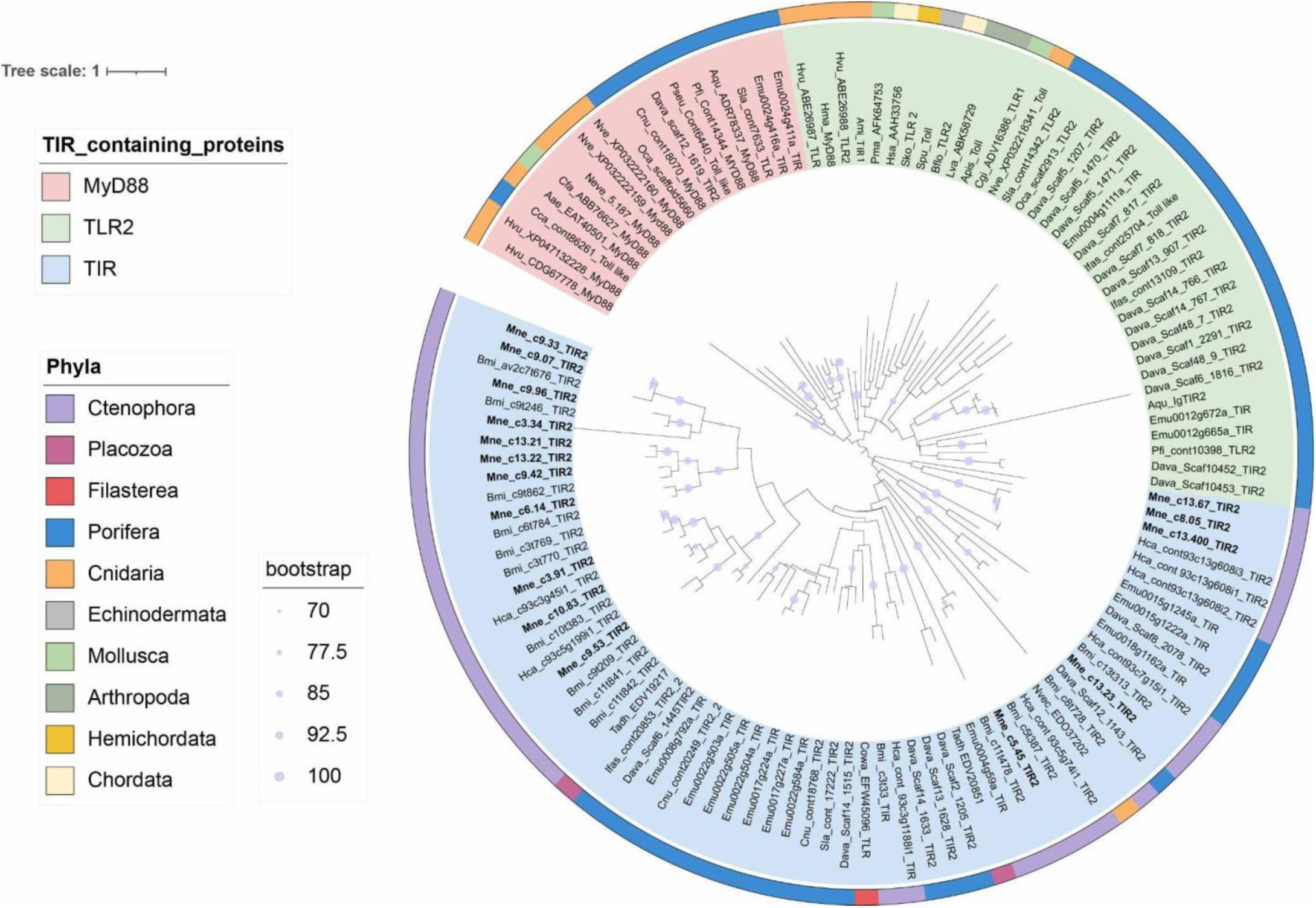
Phylogenetic analysis of TIR2 pfam domain. Phylogenetic analysis of the TIR2 domain based on GTR Bootstrap model and an estimated gamma shape parameter and 100 independent searches, generated with RAxMLv.8 (raxmlgui, v2.0.1). Sequences from TLR, Myd88 and TIR domain were extracted from NCBI for most of the phyla. Hmmer profiles were generated to extract sequences from the genomes of the ctenophores *Mnemiopsis leidyi*, *Bolinopsis microptera* and *Hormiphora caliphornensis*, and the genomes of the sponges *Dysidea avara* and *Ephydatia muelleri*.

The TIR2 containing proteins of *M. leidyi* were exclusively found in the TIR containing group, together with sequences from Filasterea, Placozoa, the two other ctenophore species (*B. microptera* and *H. californensis*) and Porifera (Figure 4). Eleven out of sixteen sequences were grouped together with other ctenophore TIR2 containing sequences, mainly of *B. microptera,* indicating the existence of unique structures of TIR2 sequences in ctenophores, while the other five sequences were spread in smaller groups together with the other ctenophore species (*B. microptera* and *H. californensis*) but also mixed with TIR sequences from other phyla. This shows first that there is an expansion of genes containing this domain in ctenophores, and second that there are also sequences with shared or similar structure with other early diverging metazoans (Figure 4).

Finally, we looked for any functional evidence among the 153 *M. leidyi* genes that contained TIR2, NACHT, SRCR or CTL domains. We assessed their changes in relative expression after exposure of *M. leidyi* with the pathogenic bacterium *Vibrio coralliilyticus* (VB) and re-exposure with the same strain after 6d (VBVB), compared with the control condition (CO), in which animals were exposed only to water. Most of the candidate genes were constitutively expressed, independently of the condition (Supplementary Table 3). While we did not detect any differential expression of these genes during the first exposure to the bacteria strain (VB), 32 genes were differentially expressed during the second bacterial exposure (VBVB *vs* CO condition, 21 % of all candidates) (Figure 5). Four genes belonged to the TIR2 domain-containing group, 13 genes to the NACHT domain, 12 genes to the SRCR domain and 4 genes to CTLD (Figure 5, Supplementary Table 4). In total, 12 genes were downregulated (7 NACHT, 4 SRCR, 2 CTLD genes) and 20 were upregulated (4 TIR2, 6 NACHT, 8 SRCR, 2 CTLD genes) in VBVB *vs* CO condition. Almost half of the NACHT (7 genes) and CTLD (2 genes) coding genes were downregulated while the other half were upregulated in VBVB *vs* CO condition. On the other hand, 8 out of 12 SRCR coding genes and all 4 TIR2 genes were upregulated upon exposure with VBVB (Figure 5, Supplementary Table 4), suggesting that they are involved in immune signalling and defence.

**Figure 5.**
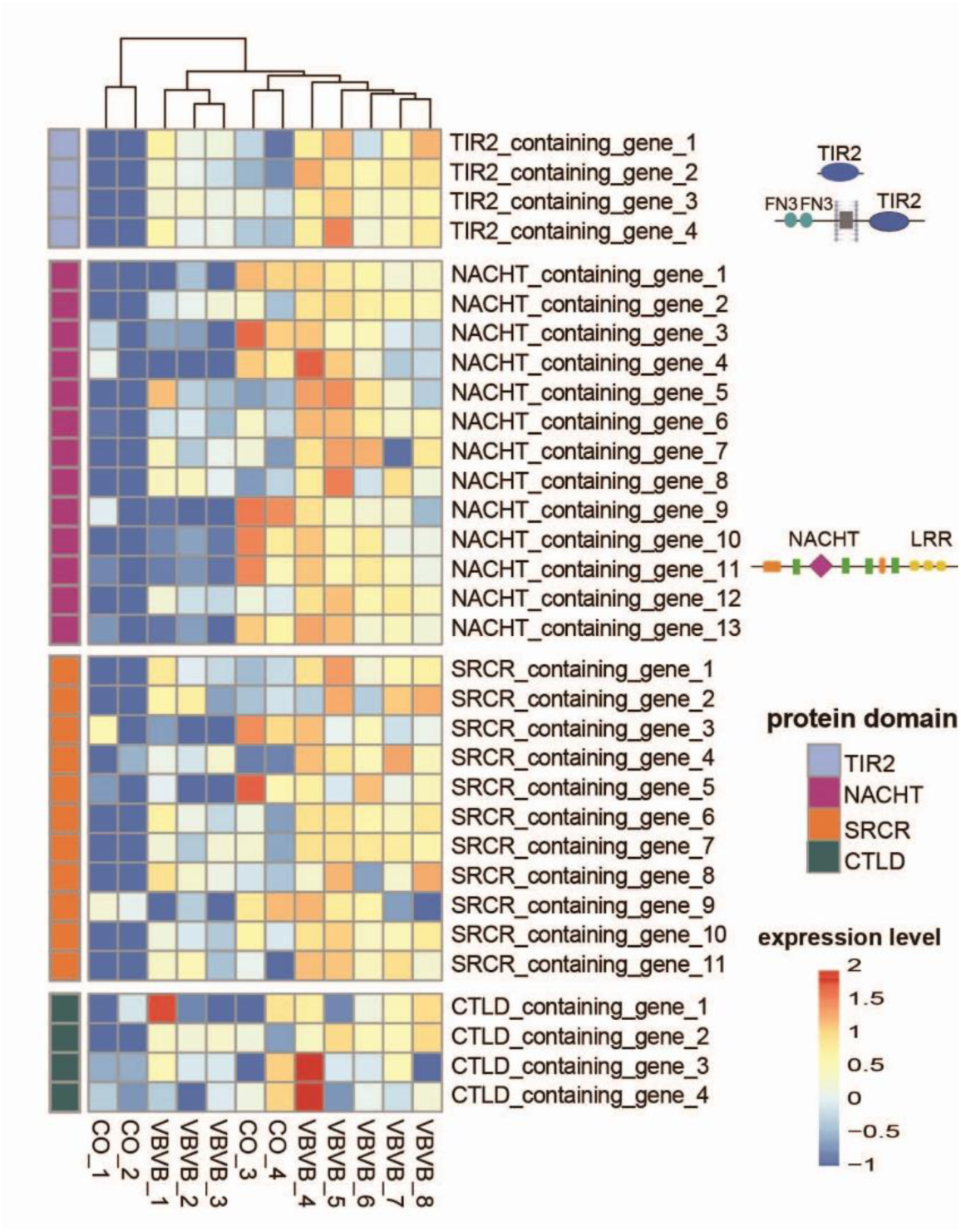
Comparative expression analysis. Heatmap with differential expression of, TIR2, NACHT, SRCR and CTLD containing genes identified in the genome of *M. leidyi* 6d after exposure with a pathogen bacterium *Vibrio coralliilyticus* (VBVB), compared to the control condition (CO). The gene expression in the heatmap increases from blue to red. The differential gene expression analysis was conducted with the edgeR package. TIR, Toll/interleukine-1; SRCR, Scavenger Receptor Cysteine-Rich; CTL, C-type Lectin.

## Discussion

We present the first chromosome-level genome for the ctenophore *Mnemiopsis leidyi*, obtaining first insights of the immune gene repertoire of this organism. The *M. leidyi* genome features almost the full diversity of conserved protein domains that are part of all other metazoan immune genes. Among the protein domains, we further studied, they all revealed large expansions and some of them had also unique architectures in *M. leidyi*. We also observed that some of the genes, containing those immune-related conserved domains, were also differentially expressed after exposure of *M. leidyi* with the pathogen bacterium *V. coralliilyticus*. This indicates that these genes do play a role in host-microbe interaction, by potentially activating the immune system of the ctenophores in order to face potential infections by pathogens.

The genome presented in this study is of high quality and as such has a great potential to be used as a resource in future studies. As a case in point, it features a higher completeness (84.7%) than the previously published *M. leidyi* draft assembly (80.8%) (Ryan et al. 2013), and also more than the two chromosome-level ctenophore assemblies (80.8% in *H. californensis* (Schultz et al. 2021) and 81.6% in *B. microptera* (Schultz et al. 2023).

Additionally, fewer metazoan BUSCO sequences were fragmented (7.8%) and the duplication rate (4.3%) is comparable to that found in the sister species *B. microptera* (4.7%) (Figure 1C). The relatively low BUSCO score of completeness rate (< 90%) in the genome of ctenophores in general can be explained by the unique phylogenetic position of ctenophores as sister group to all other metazoans (Schulz et al 2023). This range is quite common for other early diverging metazoans (e.g., Kenny et al. 2020). Finally, the completeness for the genome annotation (76.2%) was also close to the 78.2% identified in the *H. californensis* annotation (Schultz et al. 2021) and higher than the 65.5% found in the *B. microptera* annotation (Schultz et al. 2023). Finally, our results indicate, along with the other two published genomes, that the chromosome number among ctenophores is conserved with N=13 (Supplementary Figure 2). Size distribution of these 13 chromosomes in all three ctenophore species shows a constant pattern of *B. microptera* having larger homologous chromosomes than *M. leidyi*, in both cases much larger than the corresponding chromosomes of the more distantly related *H. californensis* species (Figure 1B). The species *M. leidyi* and *B. microptera* belong to the same family of *Bolinopsidae* (Ctenophora: Lobata) with *Mnemiopsis* as a paraphyletic species within the *Bolinopsis* genus (Christianson et al. 2022), while *H. californensis* belongs to Cydippids which is phylogenetically far removed from the other two species (Whelan et al. 2017). Surprisingly, very few interchromosomal translocations of orthologous genes have been identified among the three studied species (Supplementary Figure 2).

Most of the immunity-related protein domains examined in this study were present in all three ctenophore species, suggesting that they appeared in the common ancestor of the phylum Ctenophora. Notable absences were also shared, in particular one of the studied LRR domains (LRR3) and the LRCNT domain, which usually flanks LRR domains, were not detected in any of the three species. These domains are related with the NOD-like Receptors (NLRs) in vertebrates, similarly to BIR domain (Ting et al. 2008) which was lacking in *B. microptera*. However, their presence is not prerequisite in NLR formation, especially in invertebrate NLRs which do not contain any BIR domain (Lange et al. 2011; Hamada et al. 2013; Schmittmann et al. 2021). This indicates that Ctenophora already have most of the common domains present in potential immune receptors in other animals, and so we expect that these domains will execute similar function in *M. leidyi*.

All TIR domain-containing sequences identified among all three ctenophore species were more similar to the TIR2 rather than the TIR domain. The latter is a very essential component of the innate immune system as it is part of the immune TLR receptor, the IL-1R family and the Myd88 immune modulator (Bowie & O’Neill 2000; Dunne et al. 2003; Leulier & Lemaitre 2008). Although the TIR2 domain was first detected in bacteria (Newman et al. 2006), it has been found in many metazoans (Huang et al. 2008; Zhang et al. 2011), including sponges and cnidaria (Poole & Weis 2014; Riesgo et al. 2014) and is considered to be acquired through horizontal bacterial transfer like other TIR adaptor sequences (Zhang et al. 2011). Undoubtedly, ctenophores have a relatively large expansion of TIR2 containing sequences, similar to other non-bilaterian invertebrates such as Cnidaria (Poole and Weis, 2014). However, none of the TIR2 containing proteins in our study was combined with LRR domains, characteristic of canonical form TLR receptors in vertebrates (Takeda et al. 2003; Coscia et al. 2011). Traylor-Knowles et al. (2019) have previously mentioned the presence of putative TLR homologs in *M. leidyi* based on the presence of LRR domain-containing genes in the draft genome (Ryan et al. 2013); however, from our analysis, which is based on more extensive search in a complete genome assembly, we concluded that *M. leidyi* definitely lacks canonical TLR receptors. Furthermore, no TIR2 containing proteins from *M. leidyi* or the other two ctenophores, *B. microptera* and *H. californensis*, were grouped either with TLR-like (i.e., TLR homologs with non-canonical structure) or Myd88 proteins of other metazoans in our phylogenetic analysis (Figure 4). This might suggest that non-canonical TLR-like receptors and/or Myd88 are also absent in ctenophores. Despite the phylogenetic divergence of ctenophore TIR2-containing genes, 4 of those genes were upregulated upon *V. coralliilyticus* challenge (Figure 5). Interestingly, looking at the domain architecture of those 4 differentially-expressed genes, 3 of them (gene1, gene2, gene4) are exclusively TIR2 domains while in the fourth gene (gene 3) the TIR2 appears combined with Fibronectin type III (FN3) extracellular repeats. Fibronectin type III is part of the extracellular matrix and has been shown to induce the pro-inflammatory cytokines via the activation of the TLR pathway (Kelsh et al. 2014). This suggests that TIR2-containing genes are activated after MAMPs recognition, potentially activating downstream immune pathways in ctenophores.

Another interesting finding was a large expansion of NACHT domain containing proteins, combined with other domains, including LRRs but also other repeat domains such as WD40 or TPR domains. The NACHT domain is a major component of the NLRs (Proell et al. 2008; Ting et al. 2008) and based on the domain architecture of the NACHT domain-containing proteins, we suggest that *M. leidyi* has potential *bona fide* NLR receptors but it lacks the canonical NLR receptors of vertebrates. The complete canonical NLRs in vertebrates have an N-terminal with either CARD, Pyrin, Death or BIR domain, a NACHT domain and c-terminal repeats of LRR domains (Proell et al. 2008; Ting et al. 2008), while the *bona fide* NLRs are considered those with NACHT and LRR domains (Ting et al. 2008). For example, sponges and cnidarians have the complete structure of a canonical NLR receptor, such as CARD-NACHT-LRR, but also *bona fide* NLR receptors and other NACHT containing proteins (Lange et al. 2011; Hamada et al. 2013; Yuen et al. 2014; Schmittmann et al. 2021) with similar domain architecture as found here. Our finding comes in contrast with findings in the study of Yuen et al (2014), in which they did not detect any *bona fide* NLR in the draft genome of *M. leidyi*. Similarly, Moroz et al (2014) did not detect any NLR sequence in the draft genome of the ctenophore *Pleurobrachia bachei* either. On the other hand, Traylor-Knowles et al. (2019) reported the presence of NLRs as well as TLRs in ctenophores, but based only on the presence of LRRs. Interestingly, challenging *M. leidyi* with *V. coralliilyticus*, revealed the differential expression of 13 NACHT-containing genes, including the up-regulation of one *bona fide* NLR (Figure 5). Experimental evidence of NLR activation after MAMP exposure has been previously shown in the sponge *Dysidea avara* (Pita et al. 2018). Moreover, other non-canonical NLR genes were differentially expressed after LPS and flagellin stimulation, activating caspases in *Hydra* (Lange et al. 2011). Taking the fact that we find *bona fide* NLRs in ctenophores but not any *bona fide* TLR, we suggest that NLRs appeared prior to TLR receptors in the evolution of PPRs and possibly acquired an earlier role in the immune system in metazoans. Furthermore, a comparative genomic analysis by Yuen et al. 2014 showed no evidence for the existence of *bona fide* NLRs outside the Metazoa, suggesting that *bona fide* NLRs is a metazoan-specific feature with an important role in the origin and evolution of Metazoa.

A large expansion of CTLD and SRCR containing proteins was detected in *M. leidyi*. The CTL and SR receptors are very diverse and play a role in host-microbe interaction and immunity, both in vertebrates and invertebrates (Drickamer & Taylor 2015; Pees et al. 2016). Extensive studies on CTLD containing proteins and their function in immunity have been conducted in crustaceans, nematodes and insects (Pees et al. 2016). In *M. leidyi*, different combinations of domains were found in CTLD proteins, similar to those found in the sponge *Halichondria panicea* (Schmittmann et al. 2021). We also found a large range of SRs, as has been previously described in other invertebrates (Huang et al. 2008; Rast & Messier-Solek 2008; Pancer 2000; Neubauer et al. 2016; Schmittmann et al. 2021). Interestingly, most of the domain combinations found in *M. leidyi* have also been described in both Porifera and Cnidaria (Neubauer et al. 2016; Ryu et al. 2016; Schmittmann et al. 2021). For example, *M. leidyi* also features the SR families that contain only SRCR domain (SR-I family), the SR family that contained the CUB domain, the SR family with the trypsin domain, and several SR containing proteins with the IG domain and LDL domain (Neubauer et al. 2016). In contrast, we did not detect SRCR containing proteins that have a collagen or Lectin-like Oxidized Low-Density Lipoprotein Receptor 1 (LOX1) domains, characteristic of the human SRCRs, while we identified one sequence with a combined SRCR-CTL domain, which belong to the same SR family (SR-E) (Figure 3C). Finally, we detected the cluster of differentiation 36 (CD36) domain in the genome of *M. leidyi* (data not shown), which is considered a separate SR family (SR-B family) (Silverstein & Febbraio 2009). However, previous phylogenetic analysis of metazoan CD36 had shown that the sponge and ctenophore CD36 is distinct to that of cnidarians and vertebrates (Neubauer et al. 2016), possibly meaning that CD36 has a different function in ctenophores. In any case, most of SRCRs were upregulated upon pathogen re-exposure, indicating their role in pathogen recognition as well (Figure 5). Similarly, SRCR domain containing genes were up-regulated in sponges in response to MAMPs (Pita et al. 2018) and microbes (Yuen 2016). SRCR domain containing genes were also activated in response to microbes in echinoderms and molluscs (Pancer 2000; Liu et al. 2011; Furukawa et al. 2012). One of the main functions of SRCRs is to recognise molecules from microbes and activate their elimination by phagocytosis (Hentschel et al. 2012). In mammals, deactivation or malfunction of those proteins has been linked with bacterial infections (Martínez et al. 2011). We conclude that ctenophores have CTLD and SRCR containing proteins similar to those found in other early metazoans with function in immune response (Schmittmann et al. 2021).

Overall, most of the TIR2, NACHT, CTLD, and SRCR domain coding genes were expressed constitutively during our experiment, indicating their role in the maintenance of the normal physiological state of the host and microbe homeostasis (e.g., Pita et al. 2018, Schmittmann et al. 2021). However, genes that were upregulated upon exposure of the host with the pathogen *V. coralliilyticus*, can indicate their role in host-microbe interactions and in host immune defence. On the other hand, some of the NACHT, CTL, and SRCR containing genes were downregulated upon pathogen exposure in our study (Figure 5), which could mean that they are proteins related to other physiological and metabolic processes of the host, which are halted upon pathogen exposure, due to energetic costs. Another scenario could be that these genes participate in mutualistic recognition and enhancement of a healthy host-mutualist balance, and as such they are downregulated during pathogen exposure. For instance, SRs seem to also have a role in mutualism. Experimental analysis in the cnidarian *A. pallida* demonstrated that SRs increase host tolerance to its mutualistic dinoflagellate, while disfunction of these receptors elicits an immune response towards its mutualist *S. minutum* (Neubauer et al. 2016). It has also been previously suggested that a SRCR domain containing protein in the sponge *Petrosia ficiformis* potentially plays a role in the recognition of their cyanobacterial mutualists (Steindler et al. 2007). Finally, it is also plausible that the downregulated genes we detected upon the bacterial challenge are the result of the pathogen regulating host gene repression in order to facilitate their invasion and further infection.

We conclude that the chromosome-level genome provided in this study allowed us to perform an accurate and extensive search of the immune repertoire of *M. leidyi*. Ctenophores have a large variety of common protein domains participating in invertebrate immunity, demonstrating that these domains appeared in the last common ancestor and are conserved through animal evolution. Even though their architectures do not describe canonical receptors in *M. leidyi,* it seems that they already have a functional role in sensing and recognising microbial stimuli, as their expression level is regulated upon bacterial challenge. Their unique and large expansion in *M. leidyi* is consistent with its phylogenetic position as a sister group of all other metazoans in the tree of life. This study is a first step towards understanding the evolution of the immune system and the function of ancestral immune pathways of the animal tree of life. Good quality genomes are crucial to future immunological studies on how ctenophores interact with microbial counterparts in their surroundings, how they distinguish their mutualists from pathogens, and how their interaction with microbes might allow them to become such a successful invasive species worldwide.

## Material & Methods

### Specimen collection

Two individuals of *Mnemiopsis leidyi* were collected in June 2020 from animals at Kiel fjord, Germany (54.329786N, 10.151005E) (**Figure 1A**). One individual was immediately processed for DNA purification, while the other individual was flash frozen in liquid nitrogen and kept stored in -80°C until further process.

### High-molecular-weight DNA purification

High-molecular-weight DNA was extracted with the NucleoBond® HMW DNA kit (Macherey-Nagel Gmbh & Co, Germany) according to manufacturer’s instructions, with the following modifications: (1) the recommended volumes for the lysis steps was 4-fold increased to ensure the lysis of the whole individual, and (2) an extra wash step with solution H4 was added. The quality, quantity and size of the DNA extract was assessed by NanoDrop 2000c Spectrophotometer (peolab, Germany) and Qubit 2.0 (Life Technologies, Carlsband, CA).

### Library construction, genome sequencing and scaffolding

Long-read sequencing was carried out with the Pacbio Sequel II platform at IKMB sequencing facility in Kiel. A first library was prepared using the Sequel II Binding Kit 2.0 and sequenced in CLR mode (insert size 16.5 Kb). This first SMRT Cell produced an output of 124 Gb, with a subread N50 of 11.9 Kb. Circular consensus reads were generated using the *ccs* command from the pbbioconda suite of tools (https://github.com/PacificBiosciences/pbbioconda), producing a total of 4.9 Gb with mean length of 10.0 Kb. A second library was prepared using the Sequel II Binding Kit 2.2 and sequenced in HiFi mode (insert size 15.4 Kb). This second SMRT Cell produced an output of 83 Gb, with a subread N50 of 33 Kb, where the HiFi yield was 4.3 Gb with mean length of 18.4 Kb. The assembler IPA v1.8 (https://github.com/PacificBiosciences/pbipa) was run with *local* mode (including polishing, purging haplotigs and phasing) to assemble together the reads from both libraries into a *M. leidyi* draft assembly. This assembly was used as input for Hi-C sequencing described below.

### Dovetail Omni-C library preparation and sequencing

Frozen material from the second individual of *M. leidyi* was sent to Dovetail Genomics for generating Omni-C^®^ libraries. For each Dovetail Omni-C library, chromatin was fixed in place with formaldehyde in the nucleus. Fixed chromatin was digested with DNase I and then extracted, chromatin ends were repaired and ligated to a biotinylated bridge adapter followed by proximity ligation of adapter containing ends. After proximity ligation, crosslinks were reversed and the DNA purified.

Purified DNA was treated to remove biotin that was not internal to ligated fragments. Sequencing libraries were generated using NEBNext Ultra enzymes and Illumina-compatible adapters. Biotin-containing fragments were isolated using streptavidin beads before PCR enrichment of each library. The library was sequenced on an Illumina HiSeqX platform to produce ∼ 30x sequence coverage.

### Scaffolding the assembly with Omni-C HiRise

The input Pacbio draft assembly described above and all Dovetail OmniC library reads were used as input data for HiRise, a software pipeline designed specifically for using proximity ligation data to scaffold genome assemblies (Putnam et al. 2016). Dovetail OmniC library sequences were aligned to the draft input assembly using bwa (https://github.com/lh3/bwa). The separations of Dovetail OmniC read pairs mapped within draft scaffolds were analysed by HiRise to produce a likelihood model for genomic distance between read pairs, and the model was used to identify and break putative misjoins, to score prospective joins, and make joins above a threshold. Additionally, the presence of known metazoan single-copy orthologs was analysed using BUSCO v5.3.0 to assess genome completeness. These values were compared to two previously published ctenophore genomes. *H. californensis* genome was downloaded from NCBI (BioProject PRJNA576068) and *B. microptera* genome was downloaded from Dryad repository (https://doi.org/10.5061/dryad.dncjsxm47). BUSCO-based synteny was analysed using ChromSyn (Edwards et al. 2022).

### Genome annotation

The *genomeannotator* pipeline developed in-house (https://github.com/marchoeppner/genomeannotator) was used to annotate the newly generated chromosome-level genome of *M. leidyi*. This pipeline, which uses the Nextflow workflow language, performs automatic genome annotation based on ab-initio gene prediction as well as experimental and additional model hints from multiple possible sources (Di Tommaso et al. 2017). The main source of species-specific hints was RNA-seq data generated from a previous experiment. Briefly, an initial 20-30 mg of flash frozen tissue was used and the RNA was extracted with the AllPrep DNA/RNA mini kit Qiagen, according to the manufacturer’s guidelines. The total RNA was eluted in 50 μl of RNase free water and stored at -20 °C until sequencing. The sequencing was carried out on an Illumina Hiseq platform in IKMB facilities in Kiel. This dataset is a small subset of a larger dataset (unpublished data), belonging to the RNA-seq experiment described below. A description of the RNA-seq datasets that were used for annotation can be found in Supplementary Table 6.

The *genomeannotator* pipeline was run as follows: first, RepeatModeler v2.0.4 (Smit and Hubley 2015) (was run on the genome sequence with default parameters and the option - LTRstruct. The resulting annotated consensus repeat sequences were used as input to run RepeatMasker v4.1.2 (Smit et al. 2015) as the next step. Once the genome was masked, three kinds of extrinsic hints were generated: 1) reviewed metazoan proteins were downloaded from UniProt (Bateman et al. 2017) and aligned to the masked genome using SPALN v3.0.0 (Gotoh 2008), 2) trimmed raw RNA-seq reads, as described in the previous section, were aligned to the masked genome using STAR v2.7.10a (Dobin et al. 2013), and 3) the same trimmed raw RNA-seq reads were assembled using Trinity v2.13.2 (Grabherr et al. 2011) and the resulting transcripts were aligned to the masked genome using Minimap2 v2.22 (Li 2018). All resulting alignments were used as extrinsic hints to run Augustus v3.4.0 (Stanke et al. 2008). In parallel, PASA v2.5.2 (Haas 2003) was run using the RNA-seq derived hints (2 and 3 before) to generate gene models containing UTRs and alternative isoforms. Gene models generated with Augustus and PASA were used as input to run EvidenceModeler v1.1.1 (Haas et al. 2008) to generate the final reconciled consensus set of gene models. Finally, eggNOG mapper v2.1.7 (Cantalapiedra et al. 2021) was run to functionally annotate the gene models.

### Search of specific Pfam domains related to immunity

Specific domains of the metazoan immune repertoire were selected to be searched in the chromosome-scale genome assembly of *M. leidyi* (Figure 2). First, candidate immune receptors were isolated based on their characteristic conserved domains based on PFAM ID from the automated eggNOG annotation. A further search for each domain was conducted with hmmerscan v.3.4 (http://hmmer.org/) with default parameters directly in the translated genome by using the respective PFAM database entry from the Interproscan page (https://www.ebi.ac.uk/interpro/entry/pfam/). The selected sequences were reverse blasted with Interproscan (https://www.ebi.ac.uk/interpro/result/InterProScan/) and SMART (http://smart.embl-heidelberg.de/) in order to reassure the presence of the searched domain in the respective protein sequences. A hmmerscan search for each Pfam domain of interest was also conducted in the two other published ctenophore genomes, *H. californensis* and *B. microptera*, with the same parameters, in order to identify and compare the presence or absence of these potentially immune related Pfam domains.

We further selected the proteins containing the Pfam domains related to common immune receptors (TIR2, NACHT, SRCR, CTL) to further explore their domain architecture. The structure of the domains in each protein sequence was checked with SMART in Genomic mode (http://smart.embl-heidelberg.de/) (Letunic et al. 2021). Proteins both obtained from annotation and hmmerscan were studied in each case. The relevant graph for these proteins was created with Biorender.com.

### Phylogenetic analysis of TIR2 domaining-containing proteins

We chose the TIR2 Pfam domain to estimate its divergence in *M. leidyi* compared to other phyla and understand its expansion in ctenophores. First, we retrieved different sequences that include this Pfam domain (TIR sequences, Myd88, TLR sequences) from a range of metazoans, the unicellular eukaryote species *Capsaspora owczarzaki,* and the placozoan *Trichoplax adhaerens* (Supplementary Table 5) and we performed protein alignments with MUSCLE (Edgar 2004) using the SEAVIEW software (Gouy et al. 2010). We used the alignments to construct a HMM profile for the TIR domain and we further used HMMER to extract protein sequences using this HMM profile from the genomes of *M. leidyi*, *H. californesis* (Schultz et al. 2021) and *B. microspora* (Schultz et al. 2023) and from the published genome of the sponge *Ephydatia muelleri* (Kenny et al. 2020) and the genome of *Dysidea avara* (our unpublished data).

The extracted sequences from each species were aligned consecutively to the database created above using MAFFT (Katoh & Standley 2013) in the Geneious software (Kearse et al. 2012) with default parameters. After each alignment, the consensus sequence was kept and re-blasted in SMART to verify its identity. A maximum likelihood phylogenetic tree was generated with analysis in RAxML v.8 (Stamatakis 2014) with the raxmlGUI platform v2.0.1 (Edler et al. 2021), using GTR Bootstrap model for proteins and an estimated gamma shape parameter. The node support was calculated with the algorithm via bootstrapping and 100 independent searches. The tree was further processed with figtree v.1.4.4. and the online tool iTOL (https://itol.embl.de/).

### Differential gene expression

We tested the relative expression of the genes belonging to the above categories (TIR2, NACHT, SRCR, CTLD) after two consecutive immune challenges of *M. leidyi* with the pathogen bacterium *V. coralliilyticus*. Animals reared in filtered ambient seawater were transferred into a light and temperature-controlled chamber (18°C, 8-hour day-night cycle), and subsequently submerged in the bacterial solution (10^5^ cells ml^-1^). During the first challenge, samples were taken one hour and three hours post infection (VB). After a recovery period of six days without any bacterial exposure, individuals were exposed to the same bacterial strain in a second bacterial challenge (VBVB). Samples were taken again at one and three hours. As control groups (CO), individuals were submerged in filtered (0.22µm) seawater without introduction of bacteria, and animal samples were taken at the same time points. RNA-sequencing was conducted with the Illumina Hiseq platform and the resulting transcripts were mapped to the current genome. Differential gene expression analysis was conducted with the edgeR package (Robinson et al. 2009; McCarthy et al. 2012), and genes presenting FDR ≤ 0.01 and 2-fold change differences were identified as differentially expressed. The heatmap was generated in R v. 4.3.2.

## Declarations

### Ethics approval and consent to participate

Not applicable.

### Consent for publication

Not applicable.

## Data availability

Chromosome-level genome assembly of *M. leidyi* generated in this study is available at the European Nucleotide Archive (ENA) under the study ID PRJEB71361 and accession GCA_963919725.

## Authors’ contribution

**VK:** Data analysis, graph generation, writing of the manuscript, **MTO:** Data analysis, graph generation, writing of the manuscript, **TB:** analysis of RNA-seq experiment, editing manuscript, **JF:** sequencing, **NG:** conduction of RNA-seq experiment, **DG:** high molecular weight DNA extraction, **AMG:** troubleshooting and high molecular weight DNA extraction **RAS:** study design, funding, **LP:** study design, funding, interpretation of results, editing of the manuscript, **TBHR:** study design, funding, interpretation of results, editing of the manuscript.

## Acknowledgements

We thank Dr. Ana Riesgo Gill for her help and support in phylogenetic analysis. We also thank Prof. Ute Hentschel Humeida for her input and advice during the analysis and results interpretation.

