## Supplementary Figures for "The chromosome-level genome of the ctenophore *Mnemiopsis leidyi* A. Agassiz, 1865 reveals a unique immune gene repertoire"

### Supplementary Material

**Supplementary Figure 1. Chromosome-level assembly.** Heat maps of Omni-C vs. HiRise alignment for the genome of *Mnemipsis leidyi*.

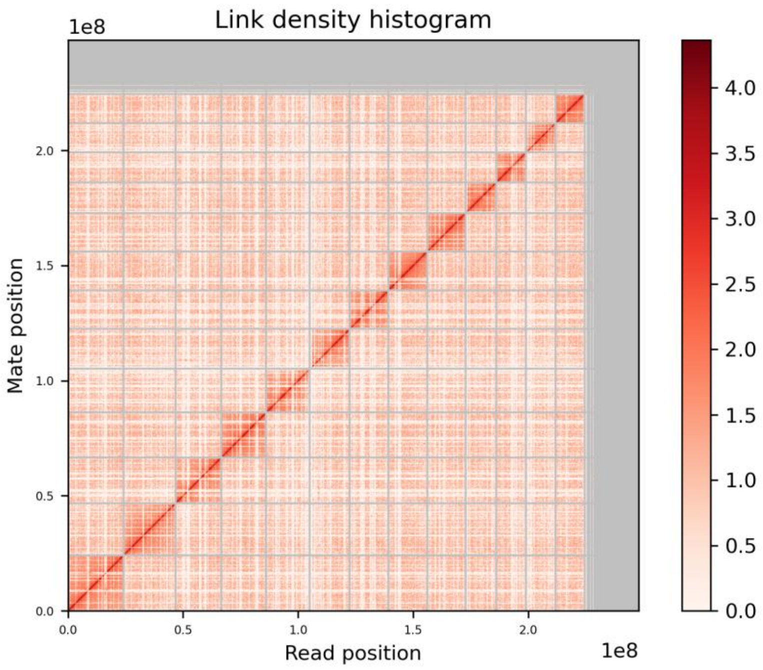


**Supplementary Figure 2. Synteny between 3 ctenophore species based on metazoan BUSCO hits.** Synteny is conserved among *Mnemipsis leidyi*, *Bolinopsis microptera* and *Hormophora californensis*. *M. leidyi* scaffolds have been relabelled accordingly: c1 [scaffold_6], c2 [scaffold_3], c3 [scaffold_2], c4 [scaffold_5], c5 [scaffold_1], c6 [scaffold_7], c7 [scaffold_11], c8 [scaffold_8], c9 [scaffold_12], c10 [scaffold_9], c11 [scaffold_4], c12 [scaffold_13], c13 [scaffold_10].

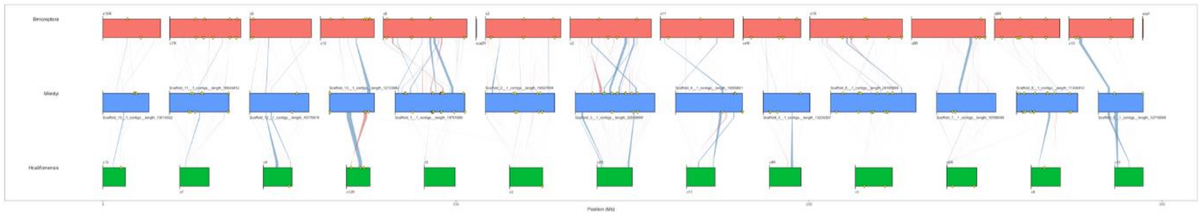
